# The mitochondrial deubiquitinase USP30 regulates AKT/mTOR signaling

**DOI:** 10.1101/2021.10.25.465793

**Authors:** Ruohan Zhang, Serra Ozgen, Hongke Luo, Judith Krigman, Yutong Zhao, Gang Xin, Nuo Sun

**Affiliations:** The Ohio State University Wexner Medical Center, Department of Physiology and Cell Biology; The Ohio State University College of Pharmacy, Division of Pharmaceutics & Pharmacology; The Ohio State University Comprehensive Cancer Center, Pelotonia Institute for Immuno-Oncology

**Keywords:** USP30, Parkin, Mitophagy, AKT, mTOR, Apoptosis, Neurogenesis, Cancer

## Abstract

Mitophagy is an intracellular mechanism to maintain mitochondrial health by removing dysfunctional mitochondria. The E3 ligase Parkin ubiquitinates the membrane proteins on targeted mitochondria to initiate mitophagy, and USP30 antagonizes this Parkin-dependent mitophagy. AKT/mTOR signaling is a master regulator of cell proliferation, differentiation, survival, and growth. Although mounting evidence showed mitophagy and AKT/mTOR signaling interact with each other during mitophagy, the specific mechanisms between Parkin/USP30 and AKT/mTOR signaling have not been elucidated. This research artificially expressed Parkin and USP30 in Hela cells and compared AKT/mTOR and apoptosis signals between Hela cells, HeLa Parkin cells, and Hela Parkin USP30 cells during mitophagy. The study’s results suggest that Parkin promotes AKT degradation via ubiquitination, which induces cell apoptosis during mitochondrial stress. On the contrary, USP30 protects AKT via deubiquitination. These findings provide new insights into Parkin and USP30’s role in cell apoptosis and physiological and pathological functions of USP30 beyond mitophagy.

## Introduction

Mitophagy, also known as mitochondria-specific autophagy, is an evolutionarily conserved cellular mechanism to recycle specific mitochondria, by being encapsulated by the structurally double membrane autophagosome [1-4]. Mitophagy plays an essential role in maintaining mitochondrial health and metabolic reactions during cellular stress, such as hypoxia or starvation [5, 6].

Mitophagy is highly regulated [7]. The PTEN (Phosphatase and tensin homolog)- induced kinase 1 (PINK1) and the cytosolic E3 ubiquitin ligase Parkin are important mitophagy promoters [8-10]. When mitochondria are damaged, PINK1 accumulates on the outer mitochondrial membrane (OMM) and recruits Parkin from the cytosol to ubiquitinate mitochondrial membrane proteins, including the translocase of the outer membrane subunit 20 (TOM20) [5, 6, 10-13]. Ubiquitin chains on the mitochondrial membrane tag the mitochondria and induce the organelle’s engulfment by the autophagosome [5, 8, 10, 11, 13]. Ubiquitin carboxyl-terminal hydrolase 30 (USP30) works as an essential checkpoint for mitophagy initiation [14-16]. USP30 is an OMM deubiquitinase that cleaves the Parkin-mediated ubiquitin chains to inhibit mitophagy [11, 15]. Loss of function mutations in Parkin and PINK1 create mitophagy defects that lead to the death of dopaminergic neurons and early onset f Parkinson’s disease [9, 17]. Inhibiting USP30 and overexpressing Parkin can promote mitophagy to rescue Parkinson’s symptoms in fruit flies [18, 19]. However, overexpressing Parkin or knocking down USP30 can be detrimental to cells, making them more vulnerable to mitochondrial stress and potential death. [15, 20, 21].

Another significant mitophagy regulator is AKT (Protein kinase B)/mTOR (The mechanistic target of rapamycin) signaling [22-24]. AKT/mTOR signaling is an intracellular pathway vital in regulating biogenesis and cell survival [25, 26]. Previous research indicated that AKT/mTOR signaling hyperactivation inhibits mitophagy and causes dysfunctional mitochondria to accumulate[24, 27]. AKT/mTOR signaling also interacts with PINK1 to promote cell survival under mitochondrial stress [21, 23, 28]. However, further investigation is required to understand better how Parkin and USP30 interact. Understanding how Parkin/USP30-AKT/mTOR interact may help us understand why cell apoptosis is induced by the overexpression of Parkin and the suppression of USP30 during mitochondrial stress. The AKT/mTOR pathway is mutated and hyperactive in 50-80% of human leukemia cases [29, 30]. AKT hyperactivation correlates with aggressive cancer progression and resistance to a plethora of chemotherapeutics [31]. The AKT/mTOR pathway is a popular anti-leukemia drug target, but the AKT inhibitor monotherapy failed due to lack of efficacy and drug resistance [32, 33] The clarification of the effects of USP30 on AKT/mTOR signalswill offer a new drug target for treating leukemia.

In our study, Hela cells, which have low Parkin expression, were transfected with Parkin (described as Hela Parkin cells). Then the Hela Parkin cells were transfected once more with Myc-USP30 to elicit an overexpression of USP30 (described as Hela Parkin USP30 cells) [16]. The cells were treated with Antimycin and Oligomycin (AO) to induce rapid PINK1/Parkin mediated mitophagy *in vitro* [10, 13, 34, 35]. The experiment’s results indicated that not only could Parkin induce mitophagy, but also promote AKT degradation during AO treatment to facilitate cell apoptosis. USP30 overexpression prevented AKT degradation during AO treatment and promoted AKT hyperactivation. Inhibiting USP30 decreased AKT in Hela Parkin USP30 cells and Jurkat T leukemia cells during mitochondrial stress and chemotherapies, promoting cell apoptosis. Further drug screening suggests that USP30 inhibitors may synergize with AKT/mTOR inhibitors in treating leukemia. Taken together, the fact that Parkin and USP30 appear to mediate AKT/mTOR signaling to regulate cell survival during mitophagy implicates USP30 as a potential drug target in leukemia treatment.

## Result

### Parkin and USP30 regulate mitophagy independent cell apoptosis during the mitochondrial stresses

We mimicked Parkin-dependent mitophagy during mitochondrial stress by treating Hela Parkin cells and Hela Parkin USP30 cells with both the mitochondrial complex III inhibitor Antimycin A and the ATP synthase inhibitor Oligomycin (AO) for up to 9 hours [36, 37]. NDP52 and OPTN are adaptor proteins that link ubiquitinated mitochondria to the autophagosome. These two proteins are degraded along with the mitochondria during mitophagy [13, 37]. As expected, Parkin induced rapid mitophagy during AO treatment, shown as the degradation of TOM20, NDP52, and OPTN in **Figure 1, a**. PARP is a universal protein that is cleaved only during apoptosis [38, 39]. During AO treatment Parkin also induced cleaved PARP during AO treatment, suggesting ongoing cell apoptosis (**Figure 1, a**). Consistent with western blot results, cell viability tests (by resazurin assay) indicated substantial cell death after 24 hours of AO treatment (**Figure 1, b**). AKT degradation and cell apoptosis all require the activation of Parkin because knocking out PINK1, the activator of Parkin, abolished both mitophagy and cell apoptosis (**Figure 1, b, and c**). Additionally, USP30 overexpression delayed mitophagy and reduced cell apoptosis during the AO treatment (**Figure 1, a, and d**). By treating cells with 10ug/ml ST-539, a specific USP30 inhibitor [16], Parkin mediated mitophagy and cell apoptosis resumed (**Figure 1, d, and e**). We next asked whether the Parkin/USP30-regulated cell apoptosis during mitochondrial stress is mitophagy dependent. Autophagy-Related-Gene 5 (ATG5) is essential for autophagosome formation. Knocking out ATG5 undermines autophagy and mitophagy [40-42]. Cell viability data showed that knocking out ATG5 did not limit cell death during mitochondrial stress (**Figure 1, f**), suggesting the Parkin/USP30-regulated cell apoptosis during mitochondrial stress is mitophagy independent. To summarize, Parkin promotes apoptosis, while USP30 antagonizes mitophagy-independent cell apoptosis during mitochondrial stress.

**Figure 1.**
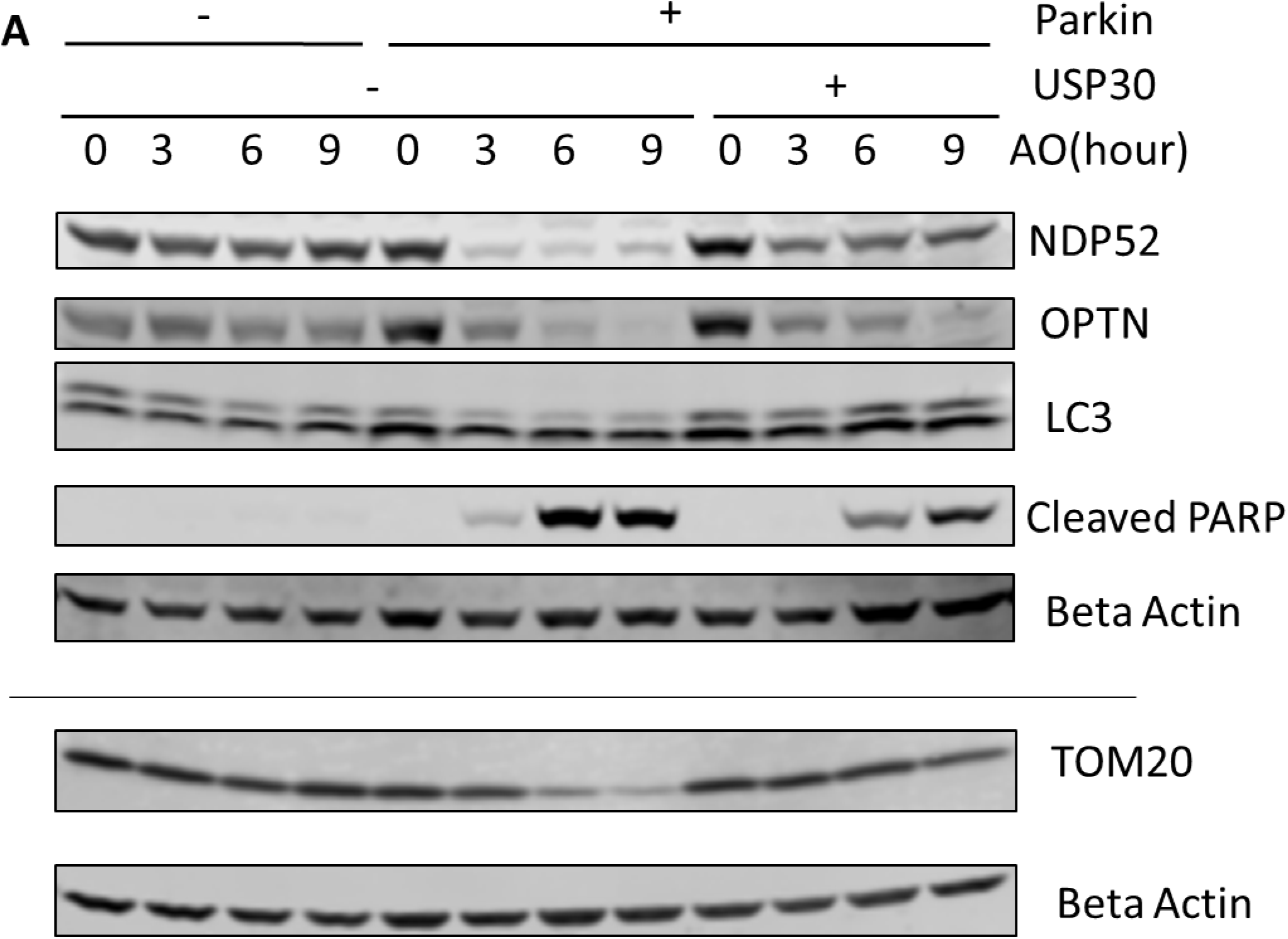
PINK1/Parkin dependent mitophagy induce apoptosis during mitochondrial stress. Figure 1a. Western blot analysis of mitophagy proteins and the pro-apoptosis signal in Hela (no Parkin) cells and Hela Parkin cells after the AO (Antimycin A + oligomycin) treatment. Cells were treated with AO at 5ug/ml for 0, 3, 6, and 9 hours. Beta Actin served as the loading control. These are representative figures from three independent experiments.

**Figure 1b.**
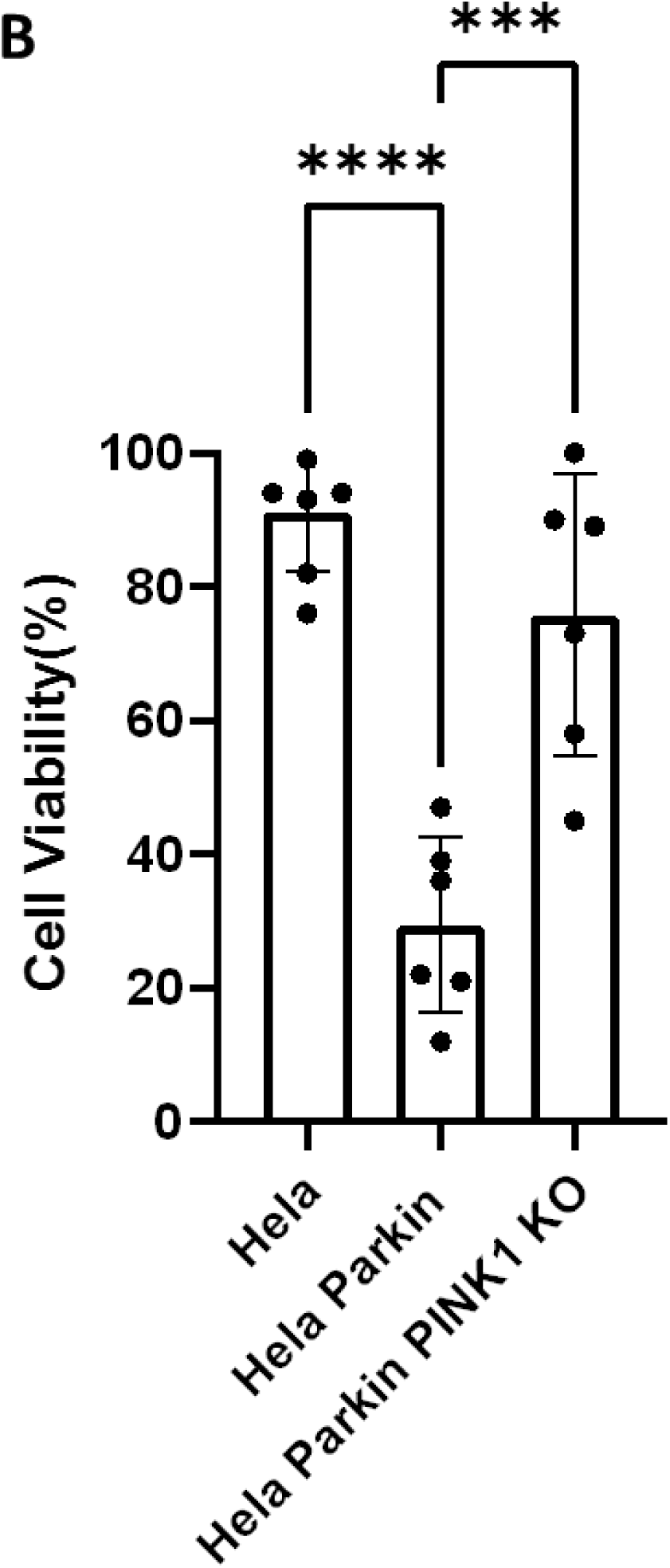
Cell viability assay of Hela (no Parkin) cells and Hela Parkin cells after AO treatment. Cells were treated with AO at 5ug/ml for 24 hours and then incubated with resazurin for 2 hours. Fluorescence was read using 544nm excitation and 590nm emission wavelength. It is a representative figure from three independent experiments.

**Figure 1 c.**
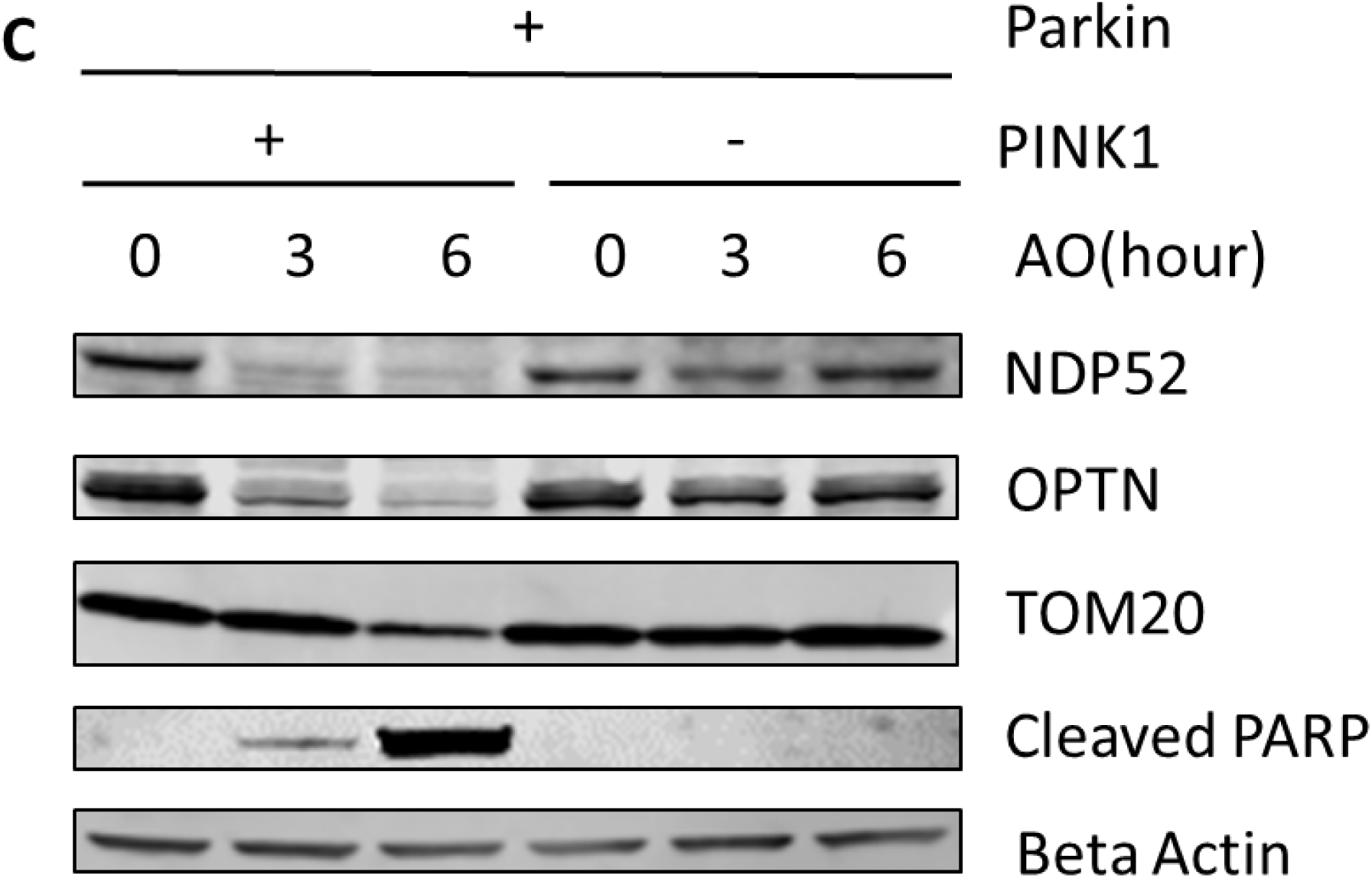
Western blot analysis of mitophagy proteins and the pro-apoptosis signal in Hela Parkin cells and Hela Parkin with PINK1 KO cells after the AO treatment. Cells were treated with AO at 5ug/ml for 0, 3, 6, and 9 hours. Beta Actin served as the loading control.

**Figure 1 d.**
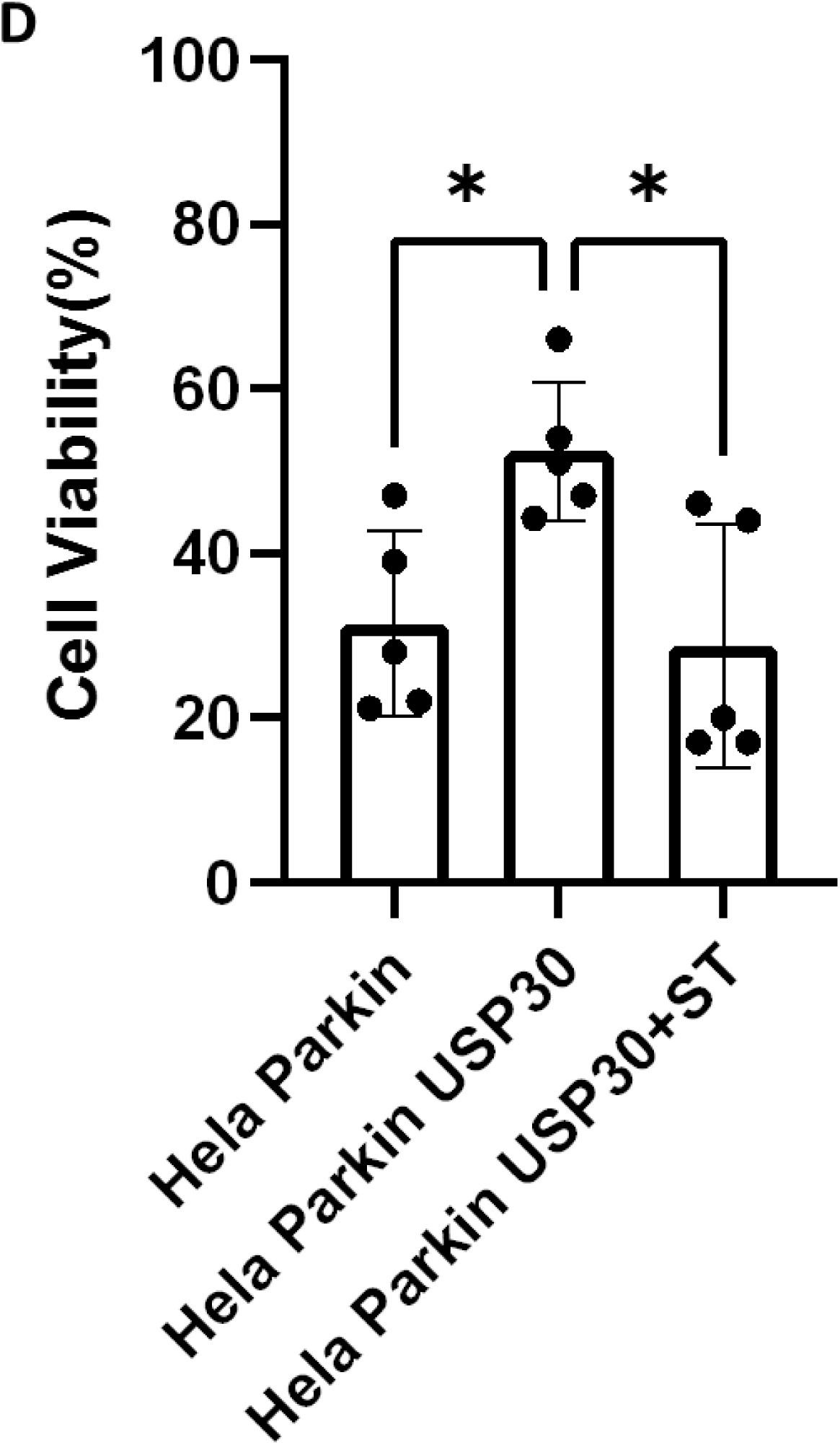
Cell viability assay of Hela Parkin cells and Hela Parkin with USP30 overexpression cells after AO treatment with or without ST-593. Cells were treated with AO at 5ug/ml w/o 10ug/ml ST-593 for 24 hours and then incubated with resazurin for 2 hours. Fluorescence was read using544nm excitation and 590nm emission wavelength.

**Figure 1 e.**
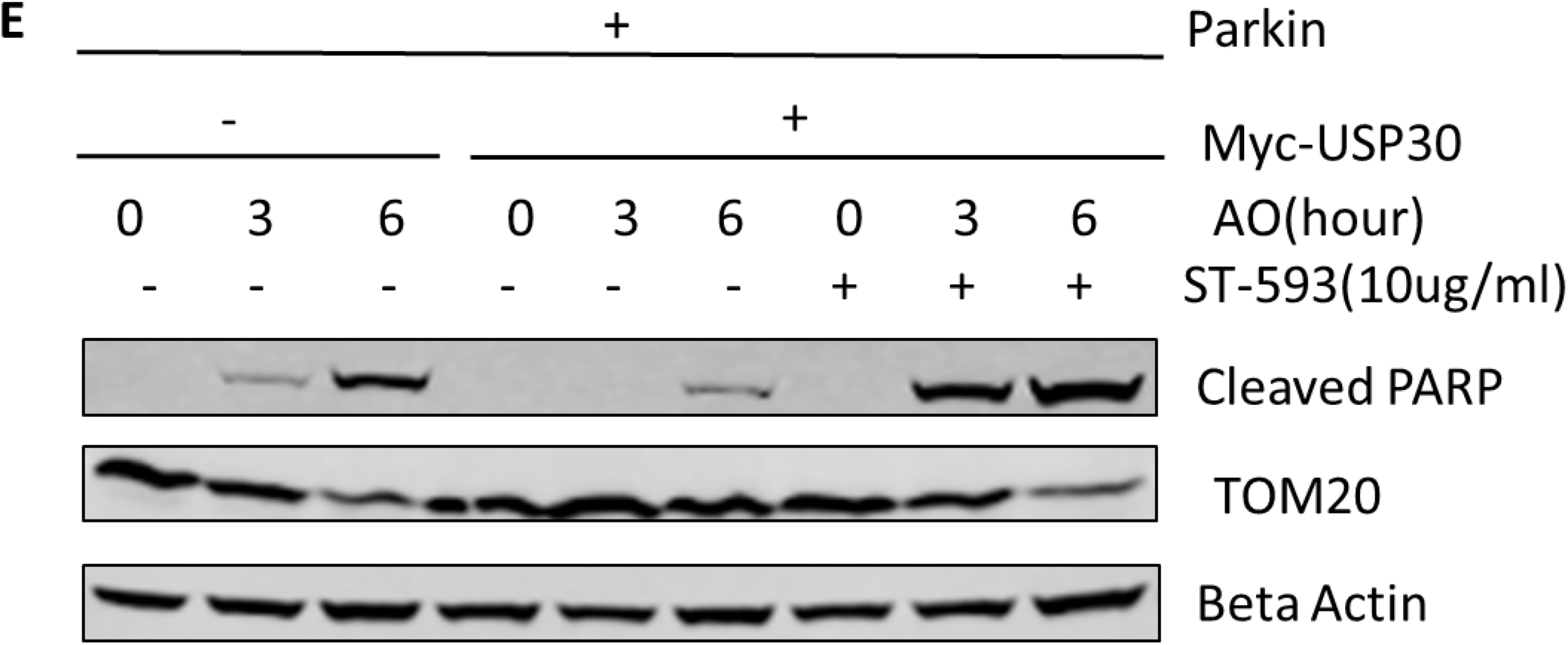
Western blot analysis of mitophagy proteins and the pro-apoptosis signal in Hela Parkin cells and Hela Parkin with USP30 overexpression cells after the AO treatment w/o ST-593. Cells were treated with AO at 5ug/ml w/o 10ug/ml ST-593 for 0, 3, and 6 hours. Beta Actin served as the loading control.

**Figure 1 f.**
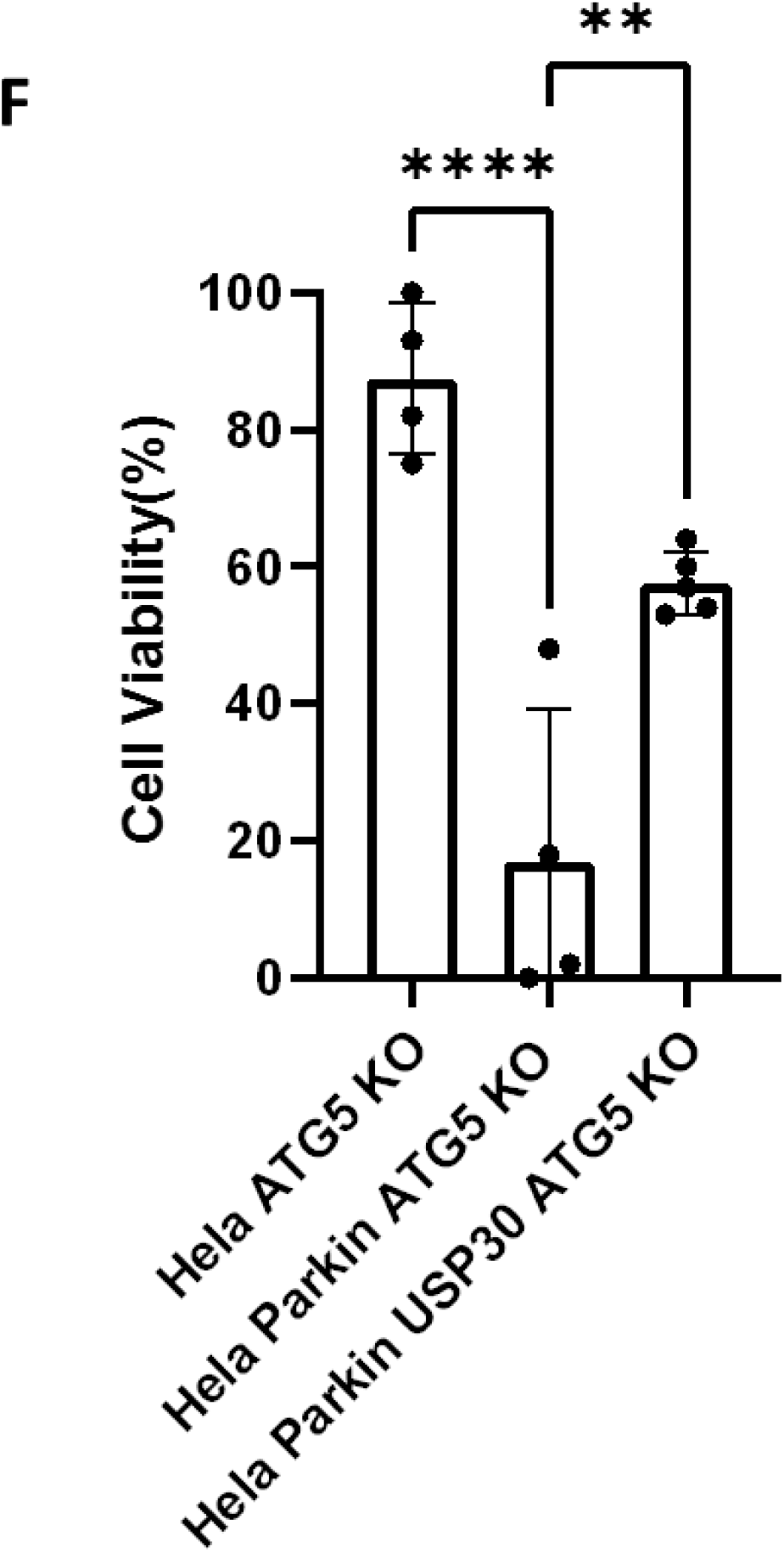
Cell viability assay of Hela ATG5 knockout cells, Hela ATG5 knockout Parkin cells and Hela ATG5 knockout Parkin USP30 cells after AO treatment. Cells were treated with AO at 5ug/ml for 24 hours and then incubated with resazurin for 2 hours. Fluorescence was read using544nm excitation and 590nm emission wavelength.

### Parkin and USP30 regulate AKT/mTOR signal which affects cell survival during mitochondrial stress

AKT/mTOR signal functions as a master signal for biogenesis and cell survival [25, 26]. Previous studies have indicated that the AKT/mTOR signal responds to multiple cellular stressors to promote cell survival [24, 25, 27]. To investigate how the AKT/mTOR signal responds to mitochondrial stress, we analyzed the substrates AKT, mTOR, and P70S6K by following three cell lines: wild-type Hela cells with minimal Parkin, our lab-created Hela Parkin cells, and our additionally transfected Hela Parkin USP30 cells during AO treatment. Western blot results indicated that AKT/mTOR signaling increased throughout AO treatment, suggesting that AKT is activated and upregulated during the induced mitochondrial stress (**Figure 2, a**). In Hela Parkin cells, AKT, mTOR, and P70S6K decreased significantly after 6 hours of AO treatment (**Figure 2, a**). This decrease suggests Parkin regulate AKT/mTOR signaling during mitophagy. USP30 overexpression significantly increased AKT’s protein level during mitophagy (**Figure 2, a**). ST-539 inhibited USP30 and decreased AKT protein levels during mitophagy (**Figure 2, b**). To test if mitophagy activity changed AKT protein levels, Chloroquine, a lysosomal inhibitor, was added to inhibit mitophagy in all cells throughout the study [43, 44]. Data suggest that the change of AKT levels during AO treatment was mitophagy independent (**Figure 2, c**). We also knocked out ATG5 in Hela Parkin and Hela Parkin USP30 cells, and results suggest mitophagy independence as well(**Figure 2, d)**. These results indicate that AKT/mTOR signal was activated under the mitochondrial stress but regulated by Parkin and USP30 in a mitophagy-independent manner.

**Figure 2.**
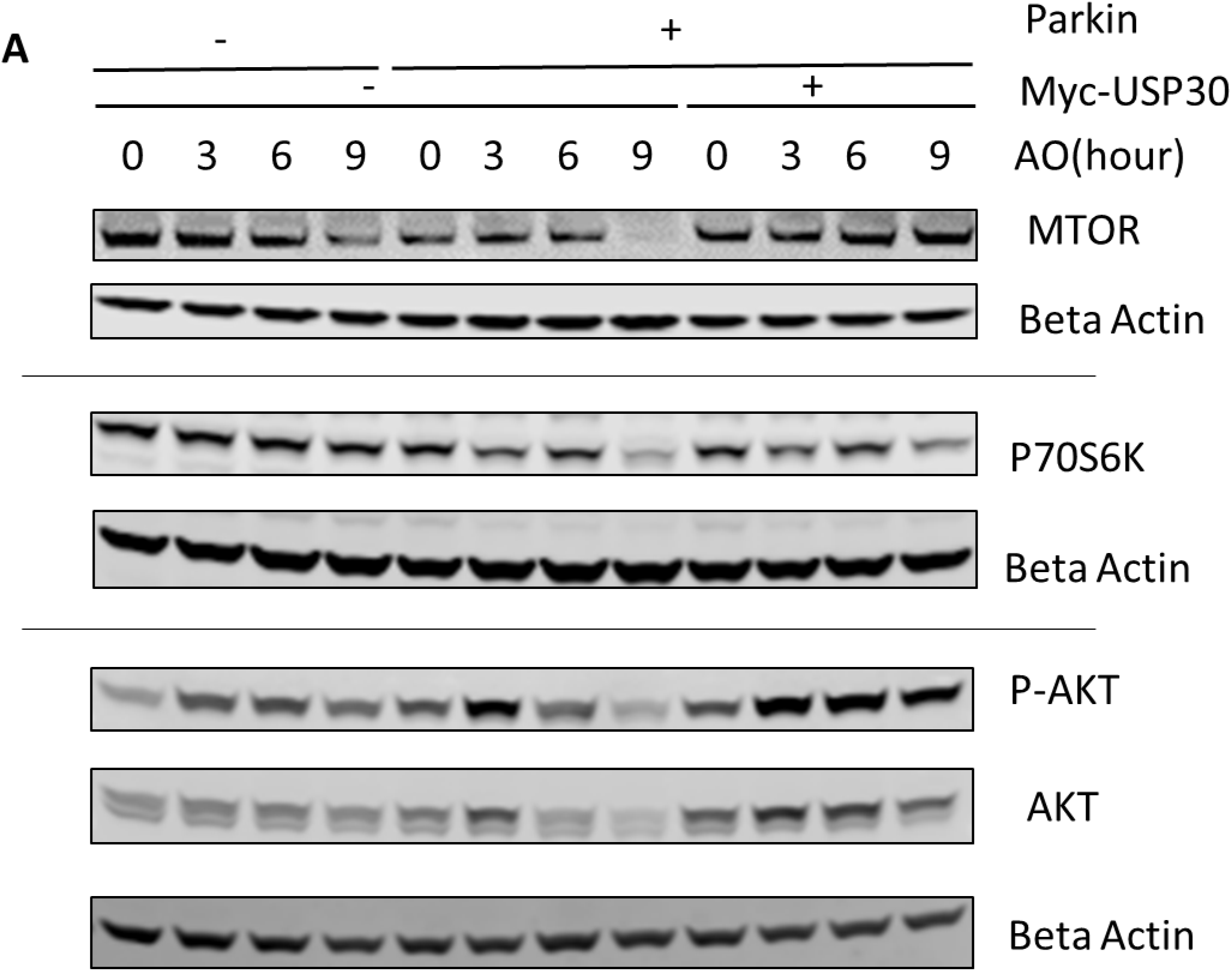
USP30 upregulates AKT/mTOR signal. Figure 2a. Western blot analysis of AKT/mTOR pathway proteins in Hela Parkin cells and Hela Parkin with USP30 overexpression cells after the AO treatment. Cells were treated with AO at 5ug/ml for 0, 3, 6, and 9 hours. Beta Actin served as the loading control.

**Figure 2 b.**
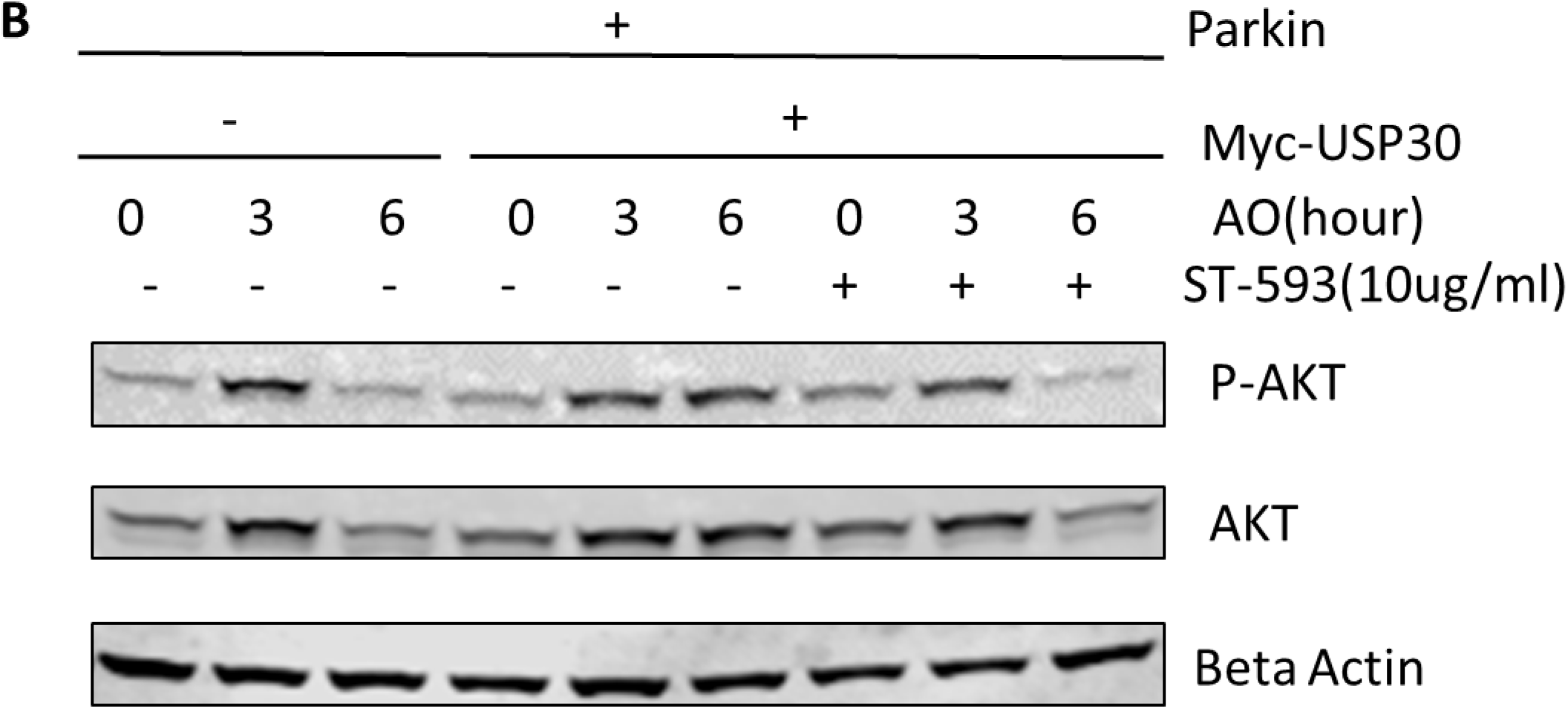
Western blot analysis of AKT signal in Hela Parkin cells and Hela Parkin with USP30 overexpression cells after the AO treatment w/o ST-593. Cells were treated with AO at 5ug/ml w/o ST-593 for 0, 3, and 6 hours. Beta Actin served as the loading control.

**Figure 2 c.**
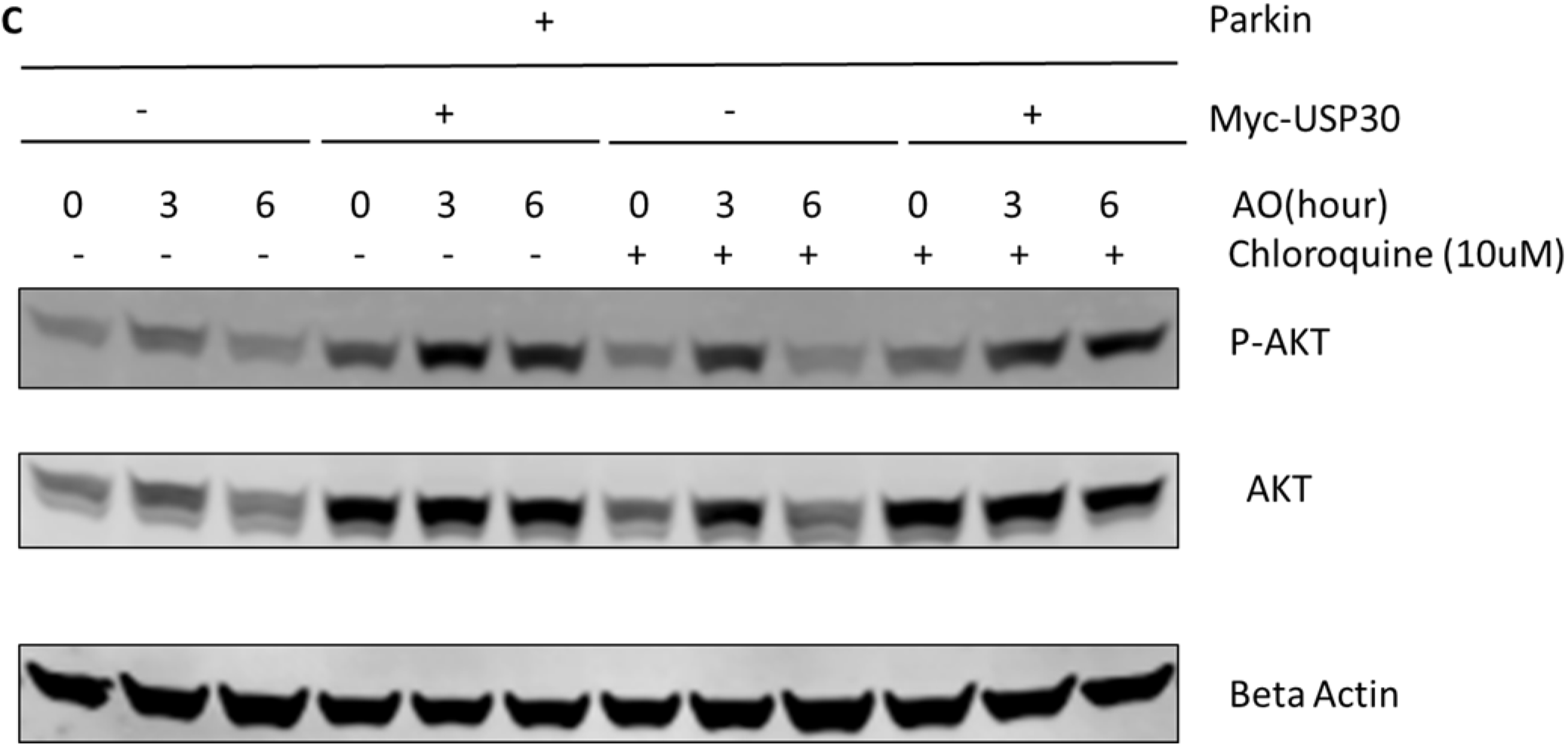
Western blot analysis of AKT/mTOR pathway proteins in Hela Parkin and Hela Parkin USP30 cells after the AO treatment w/o chloroquine. Cells were treated with AO at 5ug/ml for 0, 3, and 6 hours with DMSO or 10uM chloroquine to inhibit autophagy/mitophagy. Beta Actin served as the loading control.

**Figure 2 d.**
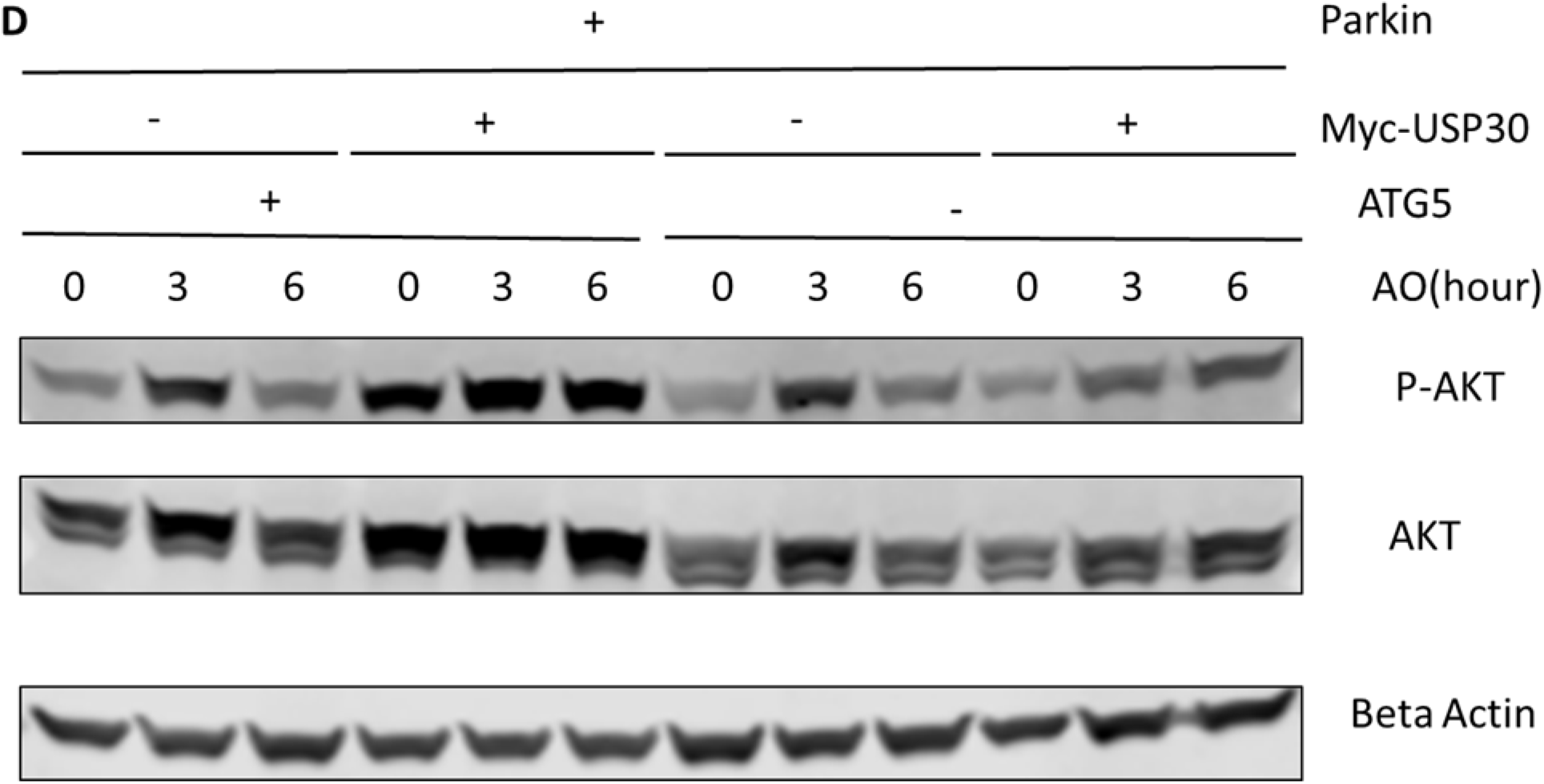
Western blot analysis of AKT/mTOR pathway proteins in Hela ATG5 knockout Parkin and Hela ATG5 knockout Parkin USP30 cells after the AO treatment. Cells were treated with AO at 5ug/ml for 0, 3, and 6 hours. Beta Actin served as the loading control.

### USP30 can be a therapeutic target in leukemia treatment

Since USP30 promotes AKT/mTOR signaling, we decided to evaluate whether USP30 could be a viable target in leukemia treatment. To start, we analyzed USP30’s effect on AKT/mTOR inhibitors by measuring the viability of Hela Parkin USP30 cells after treating them for 48 hours with MK2206 (10uM), Rapamycin (10uM), or Tronil1 (10nM) w/o ST-593. The results showed that inhibiting USP30 promoted drug-induced apoptosis significantly (**Figure 3, a**). The apoptosis in treated cells suggests that USP30 inhibitors combined with AKT/mTOR inhibitors might prove a benefit in treating leukemia. The next set of experiments focused on Jurkat cells. Jurkat cells are immortalized human T lymphocytes used to study acute T cell leukemia [45]. We evaluated Jurkat cells viability after treating them with MK2206 and ST-593.Jurkat cell growth data revealed that ST-593 is concentration-dependent, and ST-593 significantly improved MK2206’s efficacy in inhibiting cell growth. (**Figure 3, b**). Western blot results confirmed that ST-593 in conjunction with MK2006 could decrease AKT levels further and increase pro-apoptotic signaling (cleaved PARP) in Jurkat cells (**Figure 3, c**).

**Figure 3.**
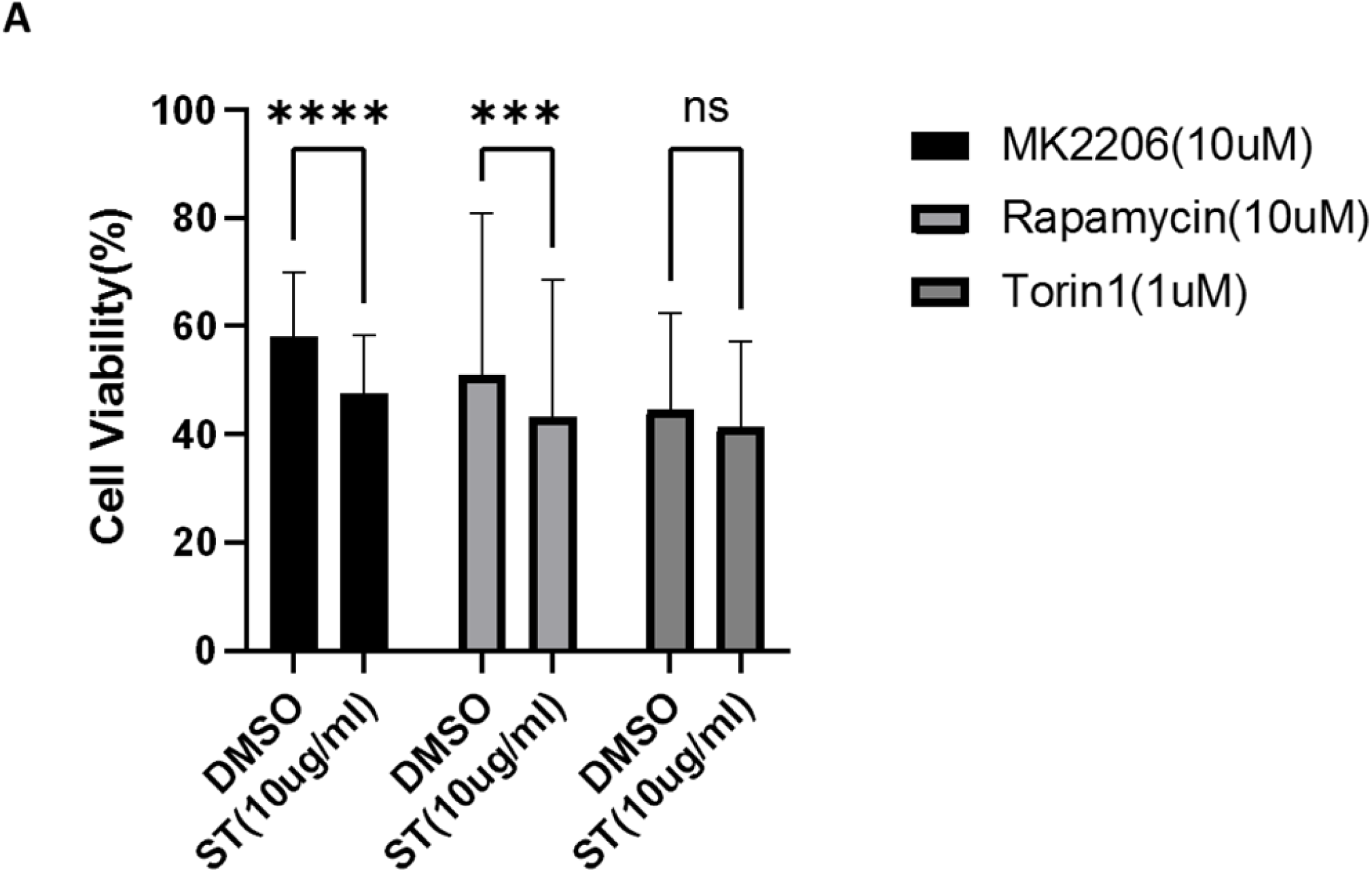
USP30 in cancer treatment. Figure 3a. Cell viability assay of Hela Parkin USP30 cells after AKT/mTOR inhibitors treatment w/o ST. Cells were treated with 10uM MK2206 or 10uM Rapamycin or 1uM Torin1 for 48hours with DMSO or 10ug/ml ST-593 and then incubated with resazurin for 2 hours. Fluorescence was read using 544nm excitation and 590nm emission wavelength.

**Figure 3 b.**
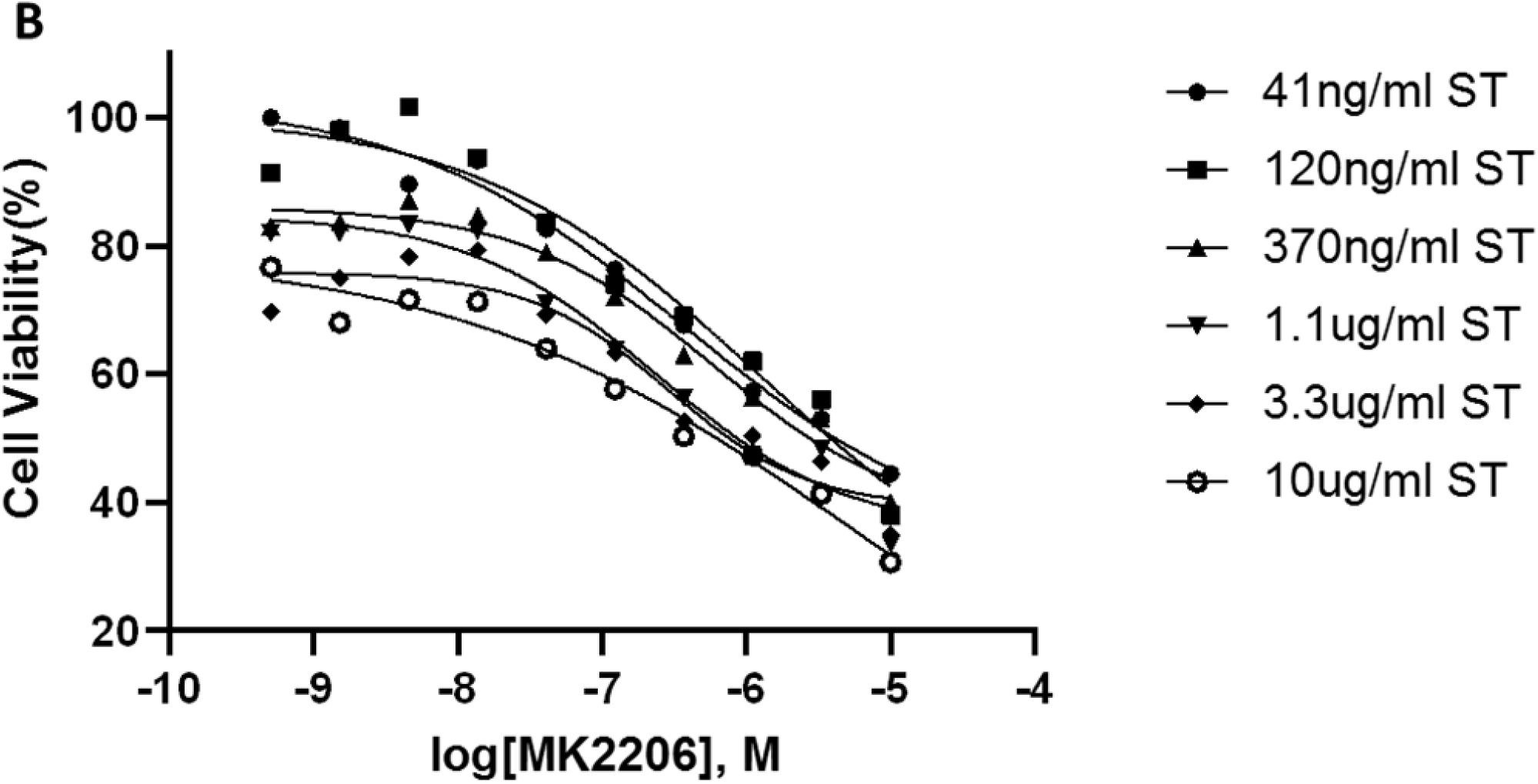
Cell viability assay on Jurkat T cells after 72 hours MK2206 treatment with ST. Cells were treated with MK2206 and ST in the concentration gradient manner for 72 hours. After the treatment, cells were incubated with resazurin for 2 hours. Fluorescence was read using544nm excitation and 590nm emission wavelength. Each dot is the mean value of three biological independent experiments. Trend lines are non-linear regression fitting curves.

**Figure 3 c.**
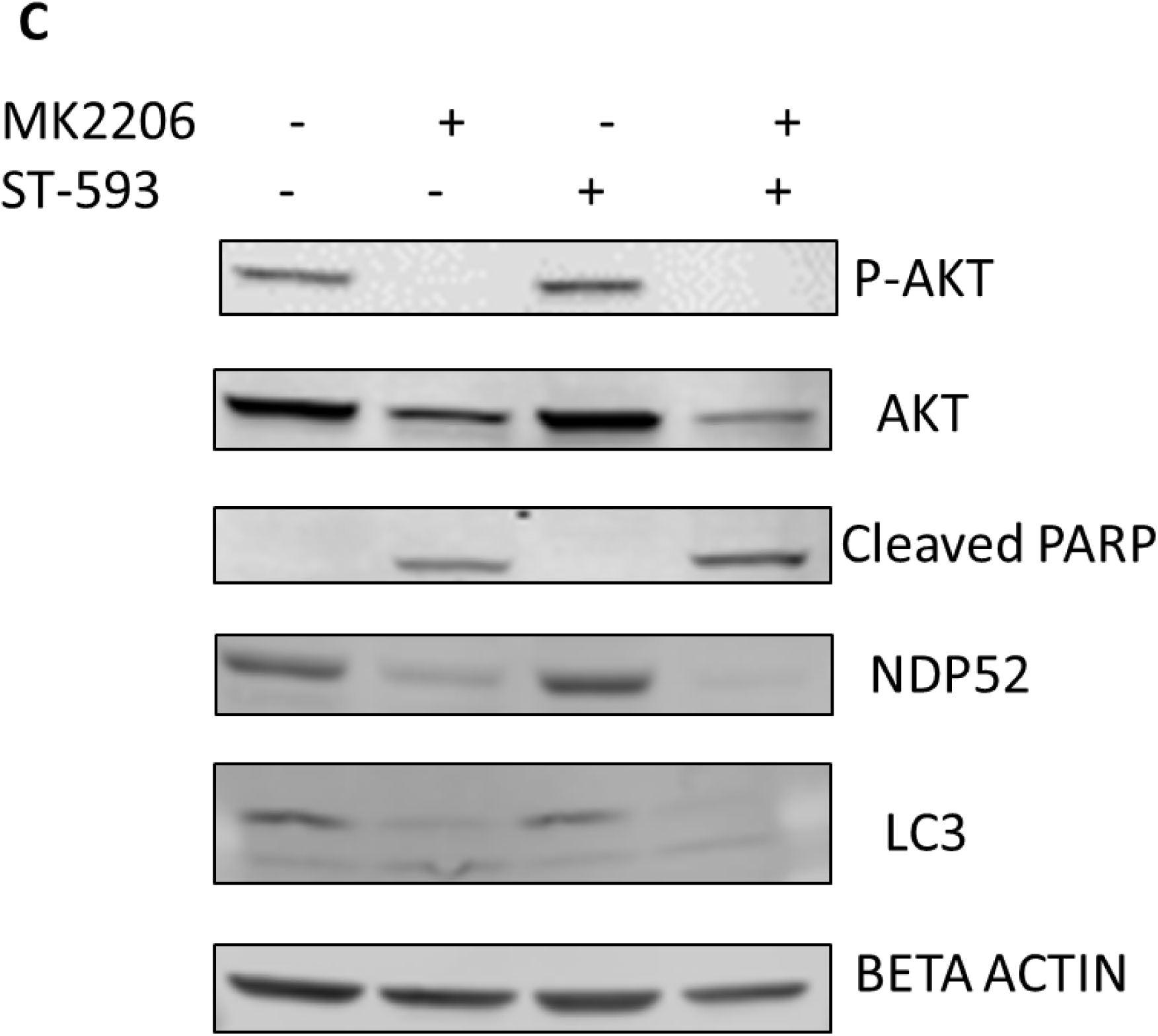
Western blot analysis of AKT, mitophagy, and pro-apoptosis signal in Jurkat T cells treated with DMSO, MK2206, or ST. Cells were treated with DMSO, MK2206, ST, or MK220+ST for 24 hours. Beta Actin served as loading control.

We used Jurkat cells to investigate the synergy between USP30 and AKT inhibitors. To do this, we reviewed an AKT/mTOR compound library composed of 322 compounds, ∼4uM with and without the compound ST-593. We evaluated Jurkat cell viability after 48 hours of treatment with various compounds. Configurations revealed that 89% of the compounds work in synergy with ST-593 to significantly suppress cell proliferation (**Figure 4, a, and b**). These results suggest a synergy exists between USP30 inhibitors, and AKT inhibitors and combining these two inhibitors to treat T cell leukemia appears promising. We ranked and reevaluated the most synergetic combinations from the previous experiment to find the most efficacious combinations (**Figure 4, c and d**). The compound, 5-lodo-indirubin-3-monoxime (Indirubin), worked best with ST-593, suppressing cell growth by 48% (**Figure 4, d**). The dose-response curve of indirubin, Glaucocalyxin A, and MK2206 w/o ST-593 showed that ST-593 promoted efficacy (**Figure 5, a-c**). In short, combining USP30 inhibitors with AKT/mTOR compounds in treating leukemia warrants further investigation.

**Figure 4.**
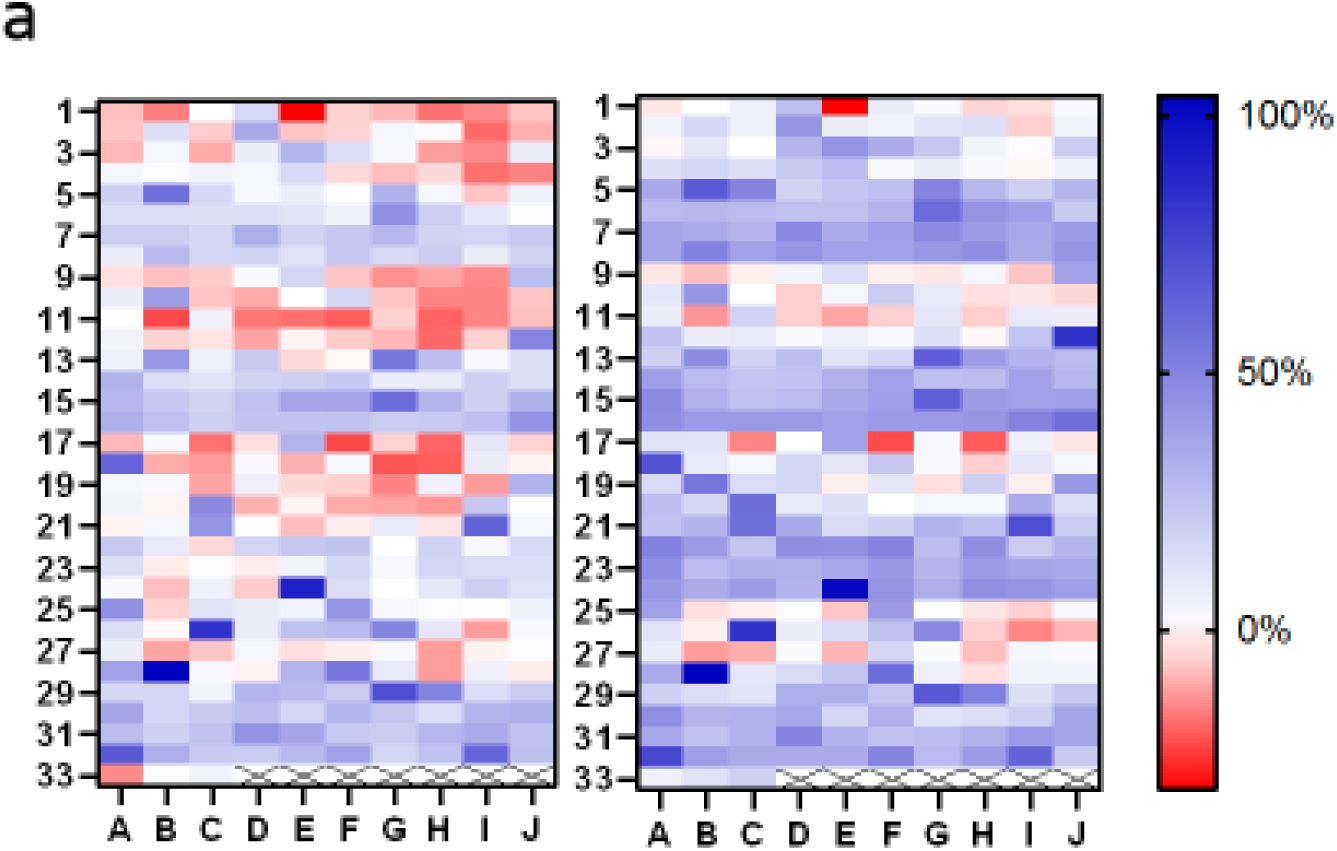
Small targeted chemical screen using the PI3K/Akt/mTOR compound library. Figure 4 a. The heat map analysis of synergy between ST-593 and AKT/mTOR inhibitors on Jurkat cells growth inhibition. Jurkat T cells were treated with the compounds (4uM) from PI3K/Akt/mTOR compound library (322 in total) w/o 10ug/ml ST-593 for 48hours, and the cell growth were analyzed by resazurin assay. The left one is the inhibition rate without ST-593, and the right one is the inhibition rate with ST-593. Data are the mean value from three biological independent experiments.

**Figure 4 b.**
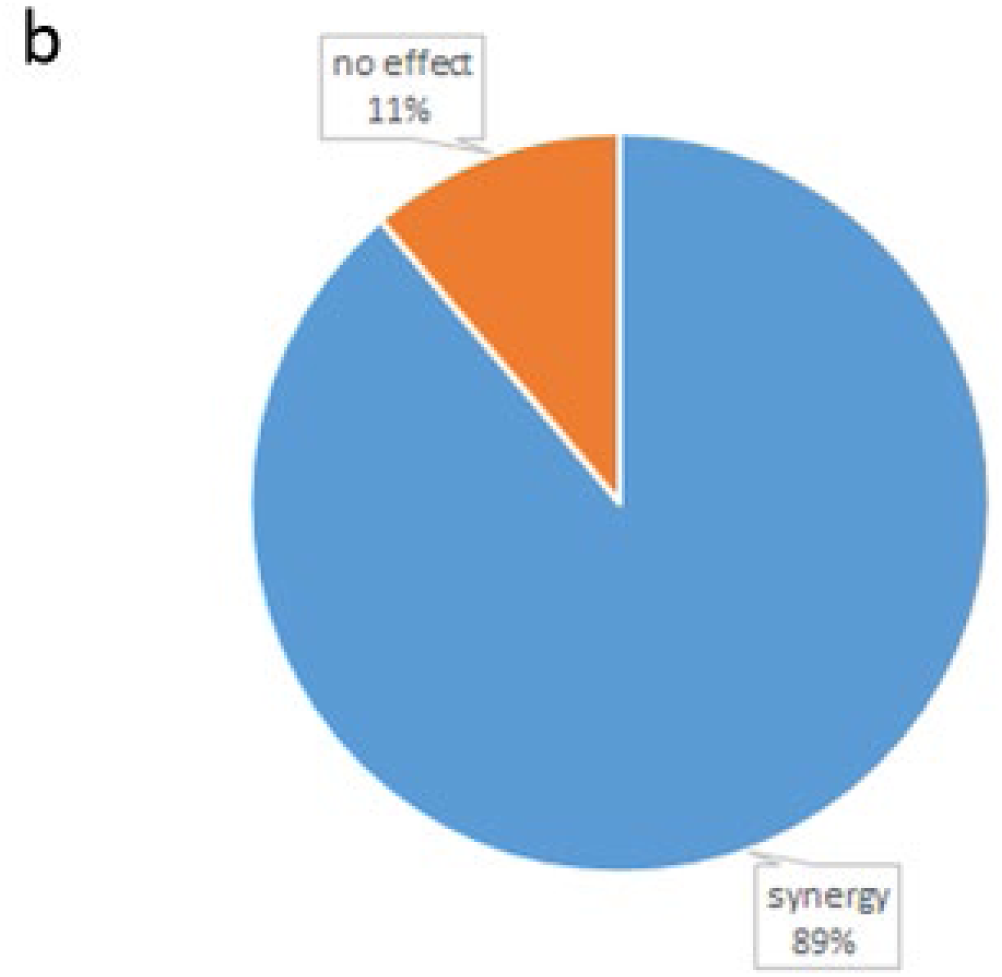
The percentage of the compounds from PI3K/Akt/mTOR compound library synergize with ST-593 on Jurkat cells growth inhibition.

**Figure 4 c.**
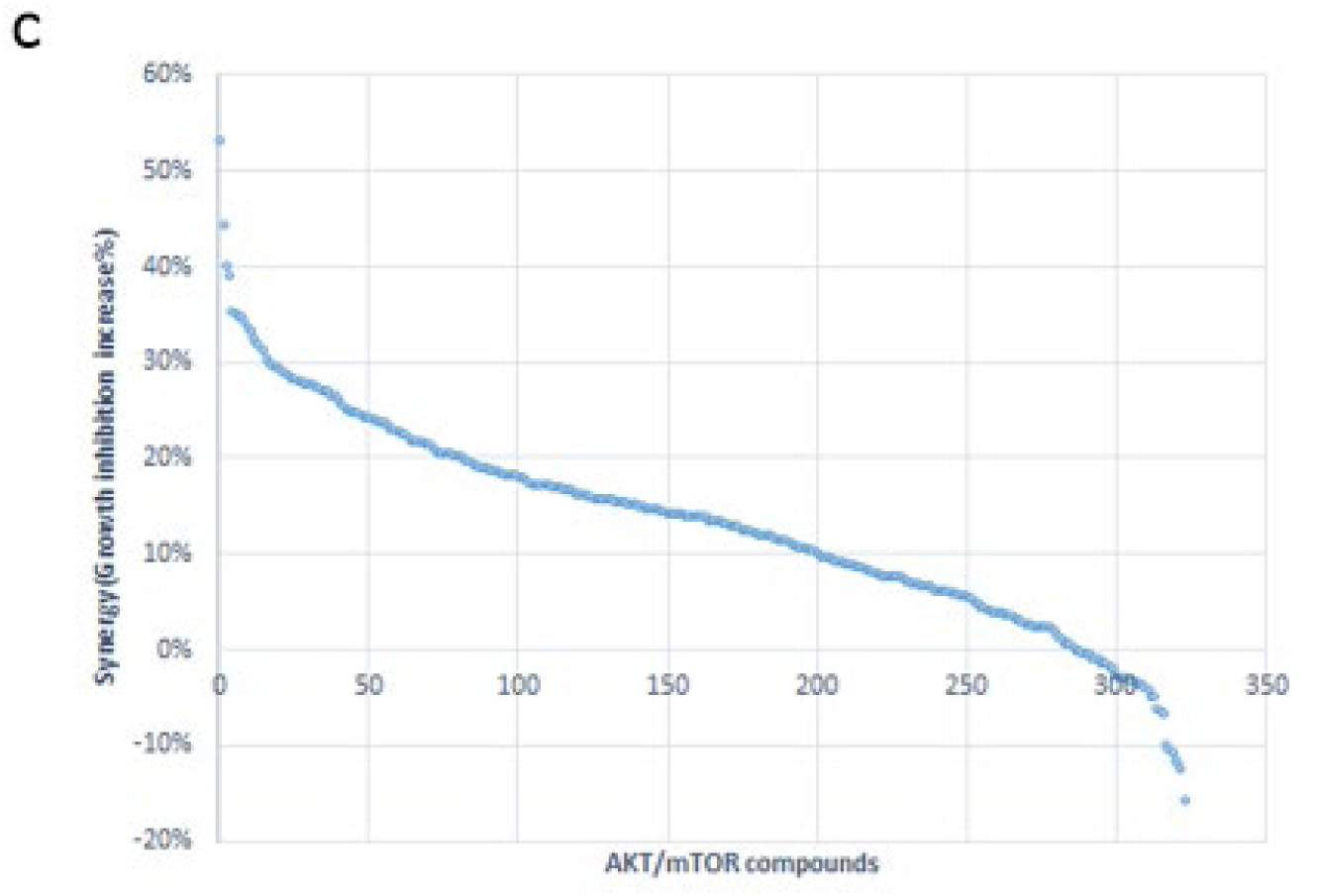
Ranks of synergistic effects between AKT/mTOR compounds and ST-593. Synergy is calculated by subtracting the growth inhibition caused by the compound alone from the growth inhibition caused by the compound and ST-593 together.

**Figure 4 d.**
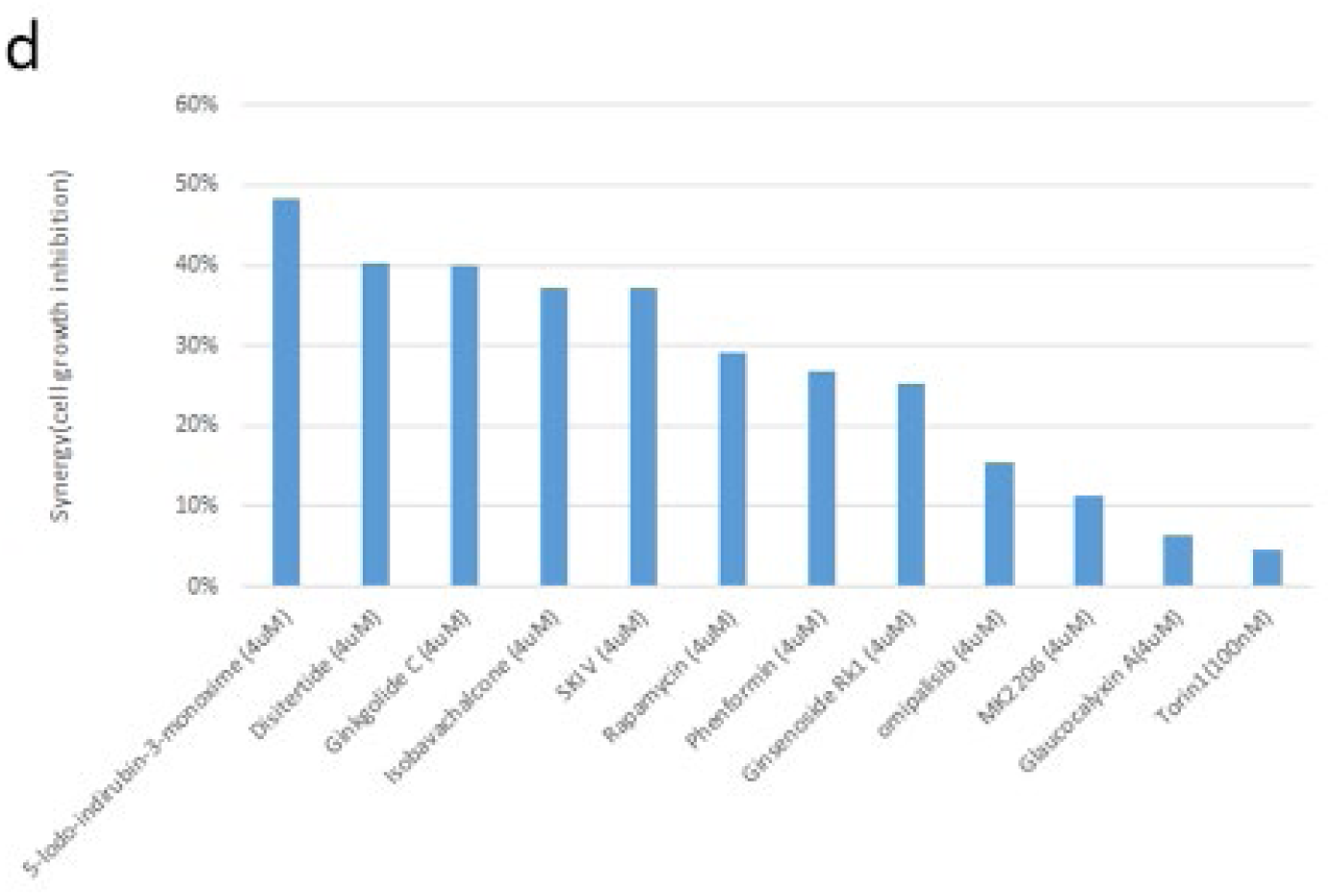
Synergy between specific compounds with ST-593. Data are the mean values from three biological independent experiments.

**Figure 5.**
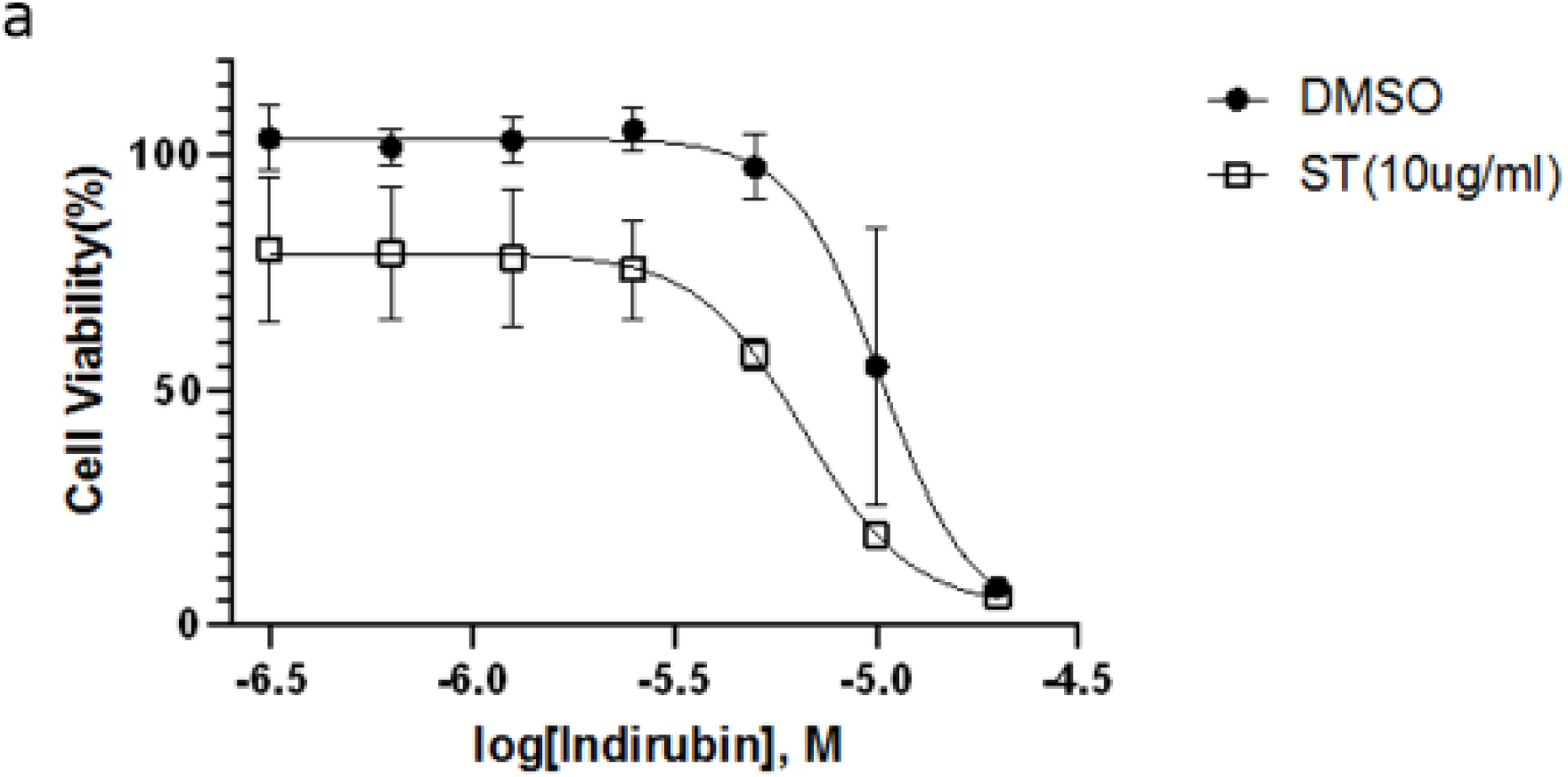
Selected inhibitors that show synergistic effects with ST-539. Figure 5 a. The dose-response curve of 5-lodo-indirubin-3-monoxime on Jurkat T cells w/o ST-593. Jurkat T cells were treated with 5-lodo-indirubin-3-monoxime in concentration gradient manner with DMSO or 10ug/ml ST-593 for 48 hours. The cell growth was analyzed using resazurin assay.

**Figure 5 b.**
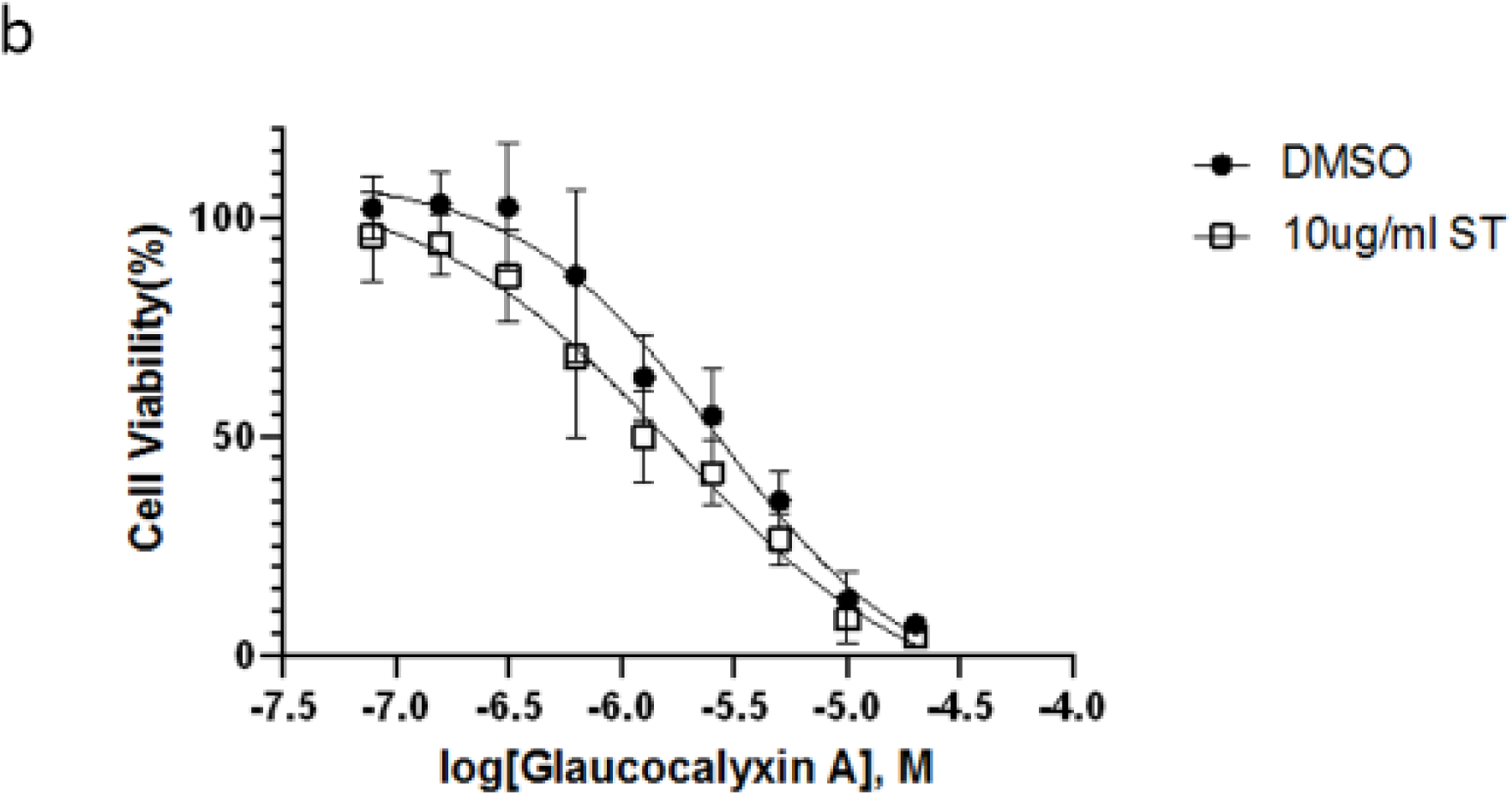
The dose-response of Glaucocalyxin A on Jurkat T cells w/o ST-593. Jurkat T cells were treated with Glaucocalyxin A in concentration gradient manner with DMSO or 10ug/ml ST-593 for 48 hours. The cell growth was analyzed using resazurin assay.

**Figure 5 c.**
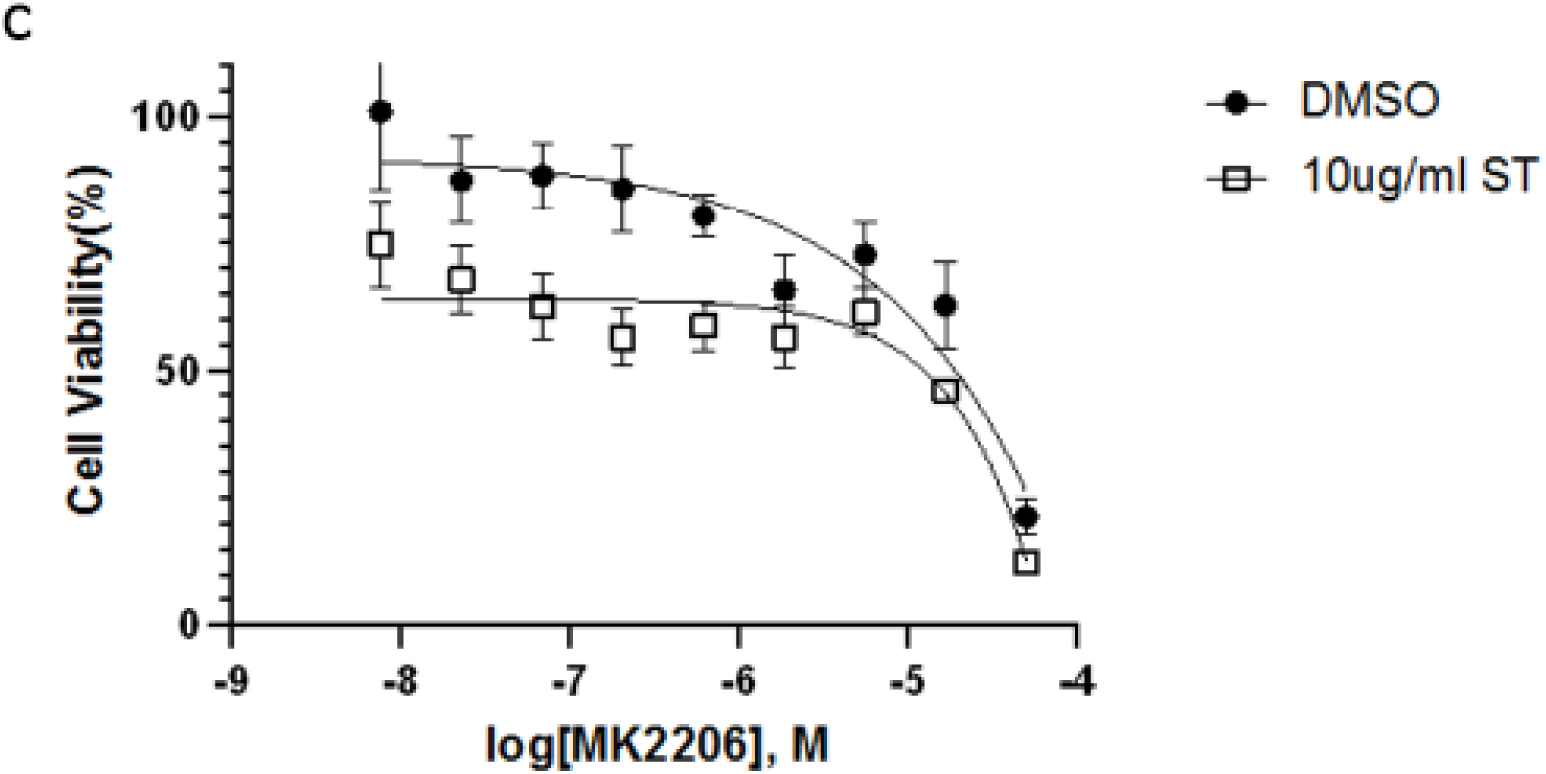
The dose-response of MK2206 on Jurkat T cells w/o ST-593. Jurkat T cells were treated with MK2206 in concentration gradient manner with DMSO or 10ug/ml ST-593 for 48 hours. The cell growth was analyzed using resazurin assay

## Discussion

Whether mitophagy promotes or works as an agonist to cancer development is unclear. Mitophagy is vital in rewiring metabolic pathways to support cancer cells’ high energy demands [46, 47]. Previous studies induced cell apoptosis by knocking down USP30 potentiated BH3, ETC inhibitors (e.g., antimycin A, oligomycin), uncouplers (e.g., FCCP), subsequently boosting Parkin-dependent mitophagy.The increase in Parkin dependent mitophagy indicates that USP30 could be a drug target in cancer treatment [15, 20]. We have demonstrated that inhibiting USP30 boosts mitophagy and downregulates AKT signaling, promoting apoptosis during mitochondrial stress. Further drug screening using Jurkat cells demonstrates that the combination of USP30 inhibitors and AKT inhibitors is efficacious in treating T cell leukemia. However, whether Parkin and USP30 regulate AKT protein levels via modulating ubiquitination during mitochondrial stress remains to be investigated.

Overall, this research revealed the connections between Parkin, USP30, and AKT signals, proving that USP30 inhibitors could be used in leukemia treatment. Future studies could focus on investigatingwhether USP30 promotes AKT signaling and drug resistance in clinical status and whether this combined therapy works on patient-derived cancer cells and murine models.

## Methods

### Cell culture and reagents

All HeLa cell lines (wild-type, HeLa Parkin, HeLa Parkin PINK1 KO, HeLa Parkin USP30, HeLa Parkin Myr-AKT, HeLa Parkin Myr-AKT K179M, HeLa Parkin AKT-T308A/S473A, HeLa Parkin AKT-T308D/473D) were grown in Dulbecco’s minimum essential medium (DMEM) with 10% fetal bovine serum (FBS) supplemented and 1% penicillin-streptomycin. HeLa Parkin and HeLa Parkin PINK1 KO cells were previously described [13]. Hela Parkin USP30 was generated using lentiviral vectors of pLVX-Puro-Myc-USP30, obtained from Addgene[48]. Jurkat T cells were grown in Roswell Park Memorial Institute Medium (RPMI-1640) with 10% fetal bovine serum (FBS) supplemented and 1% penicillin-streptomycin. For AO treatment, cells were incubated in a growth medium with 5 μM oligomycin and 5 μM antimycin A (details in figures). ST51000539 (ST-539) purchased from TimTec. Other chemicals were from Sigma-Aldrich (St. Louis).

### Western blotting

Cells were lysed in RIPA buffer (50 mM Tris-HCl, at pH 8.0; 150 mM NaCl; 1% (vol/vol) Nonidet P-40; 0.5% sodium deoxycholate, 0.1% SDS and protease inhibitor cocktail (Roche)) on ice. Primary antibodies used as described: USP30 (Sino Biological Inc., 14548-RP01, 1:500); p-AKT (Cell Signaling Technology, 4060S, 1:1000); Parkin (Cell Signaling Technology, 4211S, 1:1000); AKT (Cell Signaling Technology, 4685S, 1:1000); Cleaved PARP (Cell Signaling Technology, 5625S, 1:1000); OPTN (Proteintech, 10837-I-AP, 1:1000); p-mTOR (Cell Signaling Technology, 5536S, 1:1000); mTOR (Cell Signaling Technology, 2983S, 1:1000); TOM20 (Cell Signaling Technology, 42406S, 1:1000); NDP52 (Cell Signaling Technology, 60732S, 1:1000); p-P70S6K (Cell Signaling Technology, 9234P, 1:1000); P70S6K (Cell Signaling Technology, 9202S, 1:1000); LC3A/B (Cell Signaling Technology, 4108S, 1:1000); Beta-Actin (Cell Signaling Technology, 3700S, 1:1000). Secondary antibodies anti-rabbit (LI-COR, 926-32211, 1:15 000) and anti-mouse (LI-COR, 926-68072, 1:15 000) IgG were used to incubate membranes at room temperature for 1 hour. Images were captured using the Odyssey system (LI-Cor). One representative blot is shown of three independent experiments.

### Drug Screening

Jurkat T cells were grown in 96-wells-plates with compounds from the PI3K/Akt/mTOR compound library bought from MedChemEXpress. Cells were treated with compounds at concentrations described in the figure for 48 hours. The growth inhibition on cells was analyzed using a resazurin cell viability assay.

### Cell Viability Assay

Resazurin was bought from R&D Systems (AR002). Resazurin was added at a volume equal to 10% of the cell culture volume. Next cells with resazurin incubated for 1 to 2 hours at37 degrees. Fluorescence of the cell culture medium was read using 544nm excitation and 590nm emission wavelength.

## Acknowledge

We thank Dr. Richard Youle for generously providing us with the Hela Parkin cell lines. We thank Dr. Miyawaki for the original mt-Keima construct.

